# The association between alcohol consumption and telomere length: A meta-analysis focusing on observational studies

**DOI:** 10.1101/374280

**Authors:** Jianqiang Li, Yu Guan, Faheem Akhtar, Xi Xu, Ji-Jiang Yang, Shi Chen, Qing Wang, Hui Pan, Weiliang Qiu

## Abstract

**Background:** Both telomere length and alcohol consumption play important roles in carcinogenesis and biological age. Many efforts have been made to investigate the association between alcohol consumption and telomere length. However, no consensus has been reached yet.

**Methods:** In this article, we performed a meta-analysis to integrate the investigation results in the literature about the association between alcohol consumption and telomere length. After searching articles published between 2000 and 2016, 21 articles (including 27 analyses, total sample size 35,891) met our eligibility criteria.

**Results:** We found a significant association between alcohol consumption and telomere length (Fisher’s combined p-value = 3.52E-8 and Liptak’s weighted p-value = 8.24E-3). We also found that the significance of the association between alcohol consumption and telomere length varies with study type (cohort, case-control, or cross-sectional) and study population (Europe, Asia, American, or Australia).

**Conclusions:** Combined evidence showed that alcohol consumption is associated with telomere length. The consistent quantifications of alcohol consumption and telomere length would benefit the future aggregation of the evidence from different studies.

## Introduction

A telomere is a segment of repetitive nucleotide sequences at both tails of a chromosome, which protects the tail of the chromosome from deterioration or fusion with neighboring chromosomes[1]. Over time, the end of the chromosome becomes shorter during cell replications[1]. Once the telomere length (TL) is shortened to a critical length, the chromosomes would become unstable and the cells will immediately activate the apoptotic mechanism and lose viability[2]. So TL reflects the cell copy history and replication potential, known as the "mitotic clock" of the cell’s lifespan[1]. Defects in telomere length have been linked to several age-related diseases, premature aging syndromes, and cancers[3]. Telomere shortening may result in genomic instability during the initial stage of tumorigenesis[4].

Although telomere shortening occurs as a natural part of aging, it is known to be affected by several factors including age, gender, ethnicity, paternal age at birth, genetic mutations of telomerase, and telomere maintenance genes[5]. Telomere shortening also could be accelerated by mechanisms like oxidative stress and inflammation[5]. Furthermore, several studies suggest that psychosocial, environmental, and behavioral exposures can impact TL as well[6]. Since alcohol exposure is one of the behavioral exposures as well as a source of oxidation[7]. It would be interesting to investigate the effect of alcohol consumption (AC) on telomere length.

Alcohol is one of the most widely used recreational drugs in the world. Alcoholic drinks are classified by the International Agency for Research on Cancer (IARC) as a Group 1 carcinogen (carcinogenic to humans)[8]. IARC classifies alcoholic drink consumption as a cause of colorectal, larynx, liver, esophagus, oral cavity, pharynx, and female breast cancers; and as a probable cause of pancreatic cancer[8-10]. The World Health Organization estimates that as of 2010 there were 208 million people with alcoholism worldwide[11]. AC is the world’s third-largest risk factor for public health; in middle-income countries, which constitute almost half of the world’s population, it is the greatest risk factor for public health[12]. Also, it could reduce a person’s life expectancy by around ten years[13].

Many efforts have been made to investigate the association between AC and TL. However, no consensus has been reached yet about the AC-TL association. Some studies showed significant inverse associations between TL and alcohol use[14-16]. In other words, the more alcohol consumption, the shorter TL. However, a few studies reported significant positive associations between AC and TL. For example, Liu et al. (2009)[17] observed significantly longer TL in ever drinkers than that in never drinkers for controls of gastric cancer. Liu et al. (2011) observed significantly longer TL in ever drinkers than that in never drinkers in hepatocellular carcinoma (HCC) patients[18]. Also, several studies reported no association between AC and TL[19-30]. To facilitate the investigation of the AC-TL association, we conducted a meta-analysis.

## Methods

### Search strategy and selection criteria

This meta-analysis is based on a comprehensive and systematic search of ten research databases: ProQuest, Science Online, Wiley-Blackwell, PubMed, Science Direct, Google Scholar, Nature, Baidu scholar (http://xueshu.baidu.com/), Chinese National Knowledge Infrastructure (CNKI, https://en.wikipedia.org/wiki/CNKI, http://www.cnki.net/), and Chongqing VIP (http://lib.cqvip.com/), the last three of which are Chinese research databases. Full-text available articles in English or Chinese, published or in press between January 2000 and December 2016 were considered. Since the effect of alcohol on health is usually manifested after a long time period and clinical trials (RCTs) were usually completed in a relatively short-time period, we only focused on observational studies in this meta-analysis.

### First round of selection

The first round of selection was based on title and abstract according to the search terms (“telomere”, “alcohol”, and “ethanol”). If either the set (“telomere” and “alcohol”) or the set (“telomere” and “ethanol”) were found in title or abstract, this article would be included for further screening.

### Second round of selection

When it came to the second round of selection, only papers with full text available would be kept. The papers that did not mention any association between AC and TL were also excluded. We then grouped the remaining papers based on the types of the original studies: random clinical trial or observational study.

### Third round of selection

Since the effect of alcohol on health is usually manifested after a long time period and clinical trials (RCTs) were usually completed in a relatively short-time period, we excluded RCT studies in the third round. In the third round, we further excluded papers with low document quality.

Document quality evaluations were indispensable before any data processing, but criteria vary depending on the type of article. We applied NEWCASTLE-OTTAWA QUALITY ASSESSMENT (NOS) SCALE[31] to evaluate the quality of cohort studies and case-control studies. Meanwhile, we applied the 11-item checklist that was recommended by Agency for Healthcare Research and Quality (AHRQ) to evaluate the quality of cross-sectional studies. Based on NOS Scale, a study can be awarded a maximum of one star for each numbered item within the Selection and Outcome (for cohort studies)/Exposure (for case-control studies) categories, a maximum of two stars can be given for Comparability. The NOS ranges from zero up to nine stars[31]. According to AHRQ checklist, an item would be scored ‘0’ if it was answered ‘NO’ or ‘UNCLEAR’; if it was answered ‘YES’, then the item scored ‘1’. Article quality was assessed as follows: low quality = 0–3; moderate quality = 4–7; high quality = 8–11[32].

To pass the third round screening, we require that a cohort study or a case-control study needs to have NOS Scale ≥ 6, and a cross-sectional study need to have AHRQ ≥ 7.

Study selection was performed by YG and double-checked by JL and WQ. All disagreements were resolved through consensus by the three authors. The procedure of study selection is illustrated in **Fig 1**

**Fig 1.**
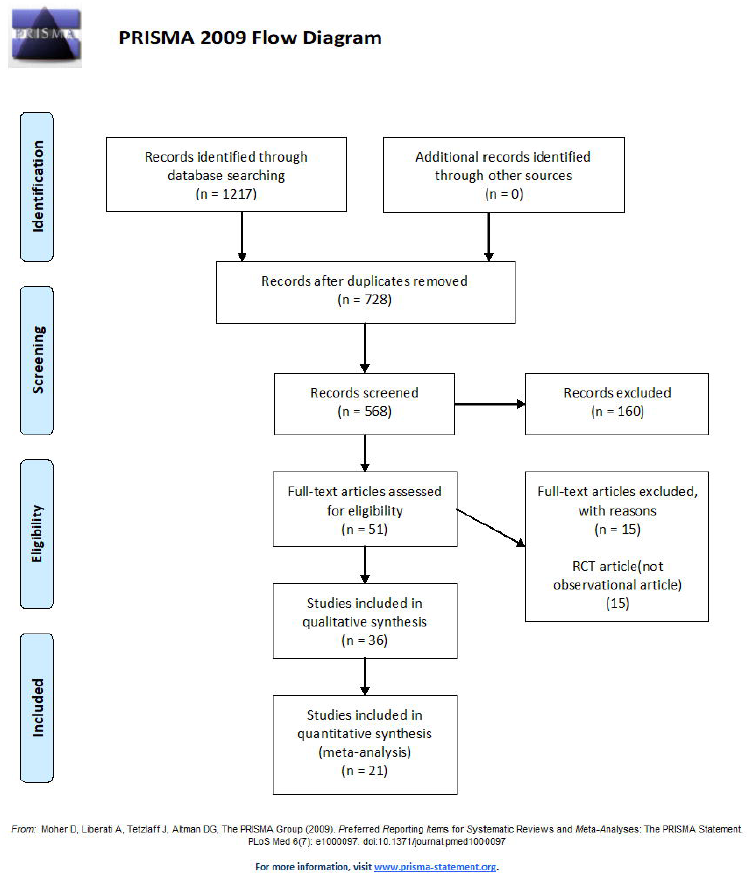
The flowchart of study selection.

### Extraction of relevant information

After the three rounds of screening, 21 articles met the requirements and were selected for further analysis. We extracted the following information relevant to the present meta-analysis: study type (case-control, cohort, or cross-sectional), significance of the association between AC and TL (test statistics and p-values), source of the population, gender, sample size, age, ethnicity, country of the participants, statistical models, quantification of TL, and quantification of AC.

### Classification of analyses

We regarded a study as a case-control study if it compared TL between two AC groups (e.g., alcohol abusers versus social drinkers). We regarded a study as a cohort study if it tested the association of AC measured at baseline to TL measured at the end of follow-up or to the change of TL from baseline to the end of follow-up. We regarded a study as a cross-sectional study if it evaluated the association of continuous-type AC to TL based on data measured at the same time period (e.g., both measured at the baseline or both measured at the end of follow-up).

### Statistical Analysis

The standard method to combine evidence from independent studies is the meta-analysis[33], in which the weighted average of the test statistics from individual studies is used to pool evidence. However, it is challenging to get appropriate test statistics for the present meta-analysis. Several different statistical models were used in the related articles to test for the association of AC to TL, such as t-test and general linear regression used in case-control studies, correlation analysis and general linear regression used in cross-sectional and cohort studies. In some studies, AC was treated as continuous variable. In other studies, AC was treated as categorical variables with different categorizations (e.g., two-category, three-category, or four-category). Even for the same number of categories, the definitions of categories might also be different. For example, Pavanello et al. (2011)[14] defined three categories as 0 – 1 drink-units/day; 2-4 drink-units/day; > 4 drink-units/day, while Houben et al. (2010)[23] defined three categories as 0 gram /day; 1-19 gram/day; >=20 gram/day. Similarly, the measurements of TL also varied among the 21 articles. Most of the studies regarded TL as a continuous variable. Houben et al. (2010)[23] and Kozlitina et al. (2012)[24] categorized TL to tertiles, Weischer et al. (2014)[34] categorized TL to quartiles, while Cassidy et al. (2010)[19] categorized TL to quintiles. Hence, the interpretations of the test statistics in different studies would be different, indicating that it is not appropriate to combine test statistics. Moreover, some studies did not provide the values of test statistics. Fortunately, almost all of the studies provided p-values and sample sizes. Therefore, we performed a meta-analysis by calculating the combined p-value as pooled evidence about the association of AC to TL. If the combined p-value < 0.05, we claim that AC is significantly associated with TL. Several methods for combining p-values have been proposed. In the present study, we used Fisher’s method. In addition, to utilize the information of the sample sizes, we used Liptak’s method[35].

Since different studies used different statistical models to investigate the association of AC to TL and did not report test statistics and their standard errors of the effect sizes, we could not directly access the potential publication bias and heterogeneity among studies. In this article, to roughly assess the heterogeneity, we used –log10(p-value) to surrogate the effect size of each study and used the inverse of sample size to surrogate the variance of the effect size. We then used –log10(Fisher’s combined p-value) and –log10(Liptak’s combined p-value), respectively, as the pooled effect size to calculated I^2^. To roughly assess the publication bias, we drew funnel plot by using signed –log10(p-value) to surrogate the effect size and using the square root of the inverse of sample size as the standard error. The sign of the effect size for a study is the same as the sign of the test statistic. If test statistic is missing, we assumed negative sign (i.e., the higher AC, the shorter TL). R package *metafor* was used to draw the funnel plot.

In this meta-analysis, we also applied Fisher’s exact test to check if the following factors affect the significance of the association between AC and TL: (1) study type (cohort study, case-control study, or cross-sectional study), (2) article goal (if testing for AC-TL association is the primary goal of the article), (3) sex-specificity (men only study, female only study, or both sex study), (4) categorization of AC (whether containing never-drinker category), and (5) study population (American, Asian, Australian, or European). For binary factors (e.g., article goal), p-values of Fisher’s exact test are obtained by directly using the hypergeometric distribution. Otherwise, the network developed by Mehta and Patel (1983, 1986)[36] and improved by Clarkson, Fan, and Joe (1993)[37] was used to calculate the p-values of Fisher’s exact test. We also applied two-sample Wilcoxon rank sum tests to check if sample size and mean age affect the AC-TL association.

A test is claimed as significant if its two-sided p-value < 0.05. All analyses were performed by using IBM SPSS Statistics Version 22.0 or R Version 3.4.2.

## Results

After the three rounds of screening, we obtained 21 articles. The information (including quality scores) of these 21 articles is listed in **Table 1**. Six of the 21 articles performed more than one analyses. There were 44 analyses in total that evaluated the association between AC and TL in the 21 articles. The information (including quality scores) of these 44 analyses is listed in **S1 Table** online. We evaluated the 27 independent analyses and discussed the remaining 17 analyses in the Discussion section. Among the 27 analyses (**Table 1**), there are four case-control studies[14, 15, 17, 18], three cohort studies[16, 34, 38], and 20 cross-sectional studies[14, 16, 19-30, 34, 38-40]. Among the remaining 17 analyses, there are ten case-control studies, three cohort studies, and four cross-sectional studies.

**Table 1.**
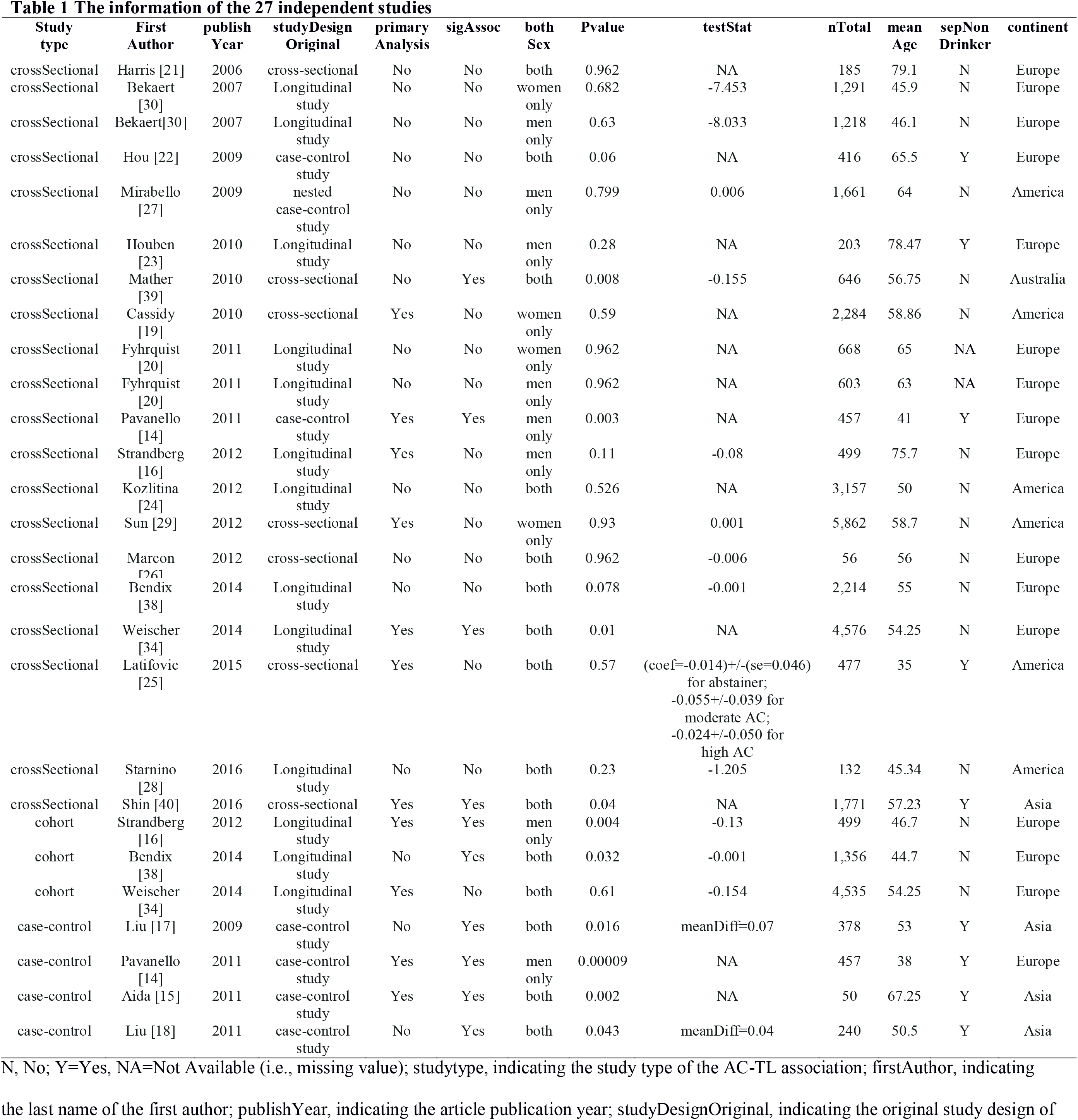
The information of the 27 independent studies.

Since different studies used different statistical models to investigate the association of AC to TL and did not report test statistics and their standard errors of the effect sizes, we could not directly access the potential publication bias and heterogeneity among studies. In this article, we used surrogate test statistics and surrogate standard errors. If we used –log10(Fisher’s combined p-value) as the pooled effect size, the value of I^2^ is equal to 34.42, indicating small to medium heterogeneity. If we used –log10(Liptak’s combined p-value) as the pooled effect size, the value of I^2^ is equal to 0, indicating small heterogeneity. **Fig 2** shows roughly symmetric funnel plot, indicating no publication bias.

**Fig 2.**
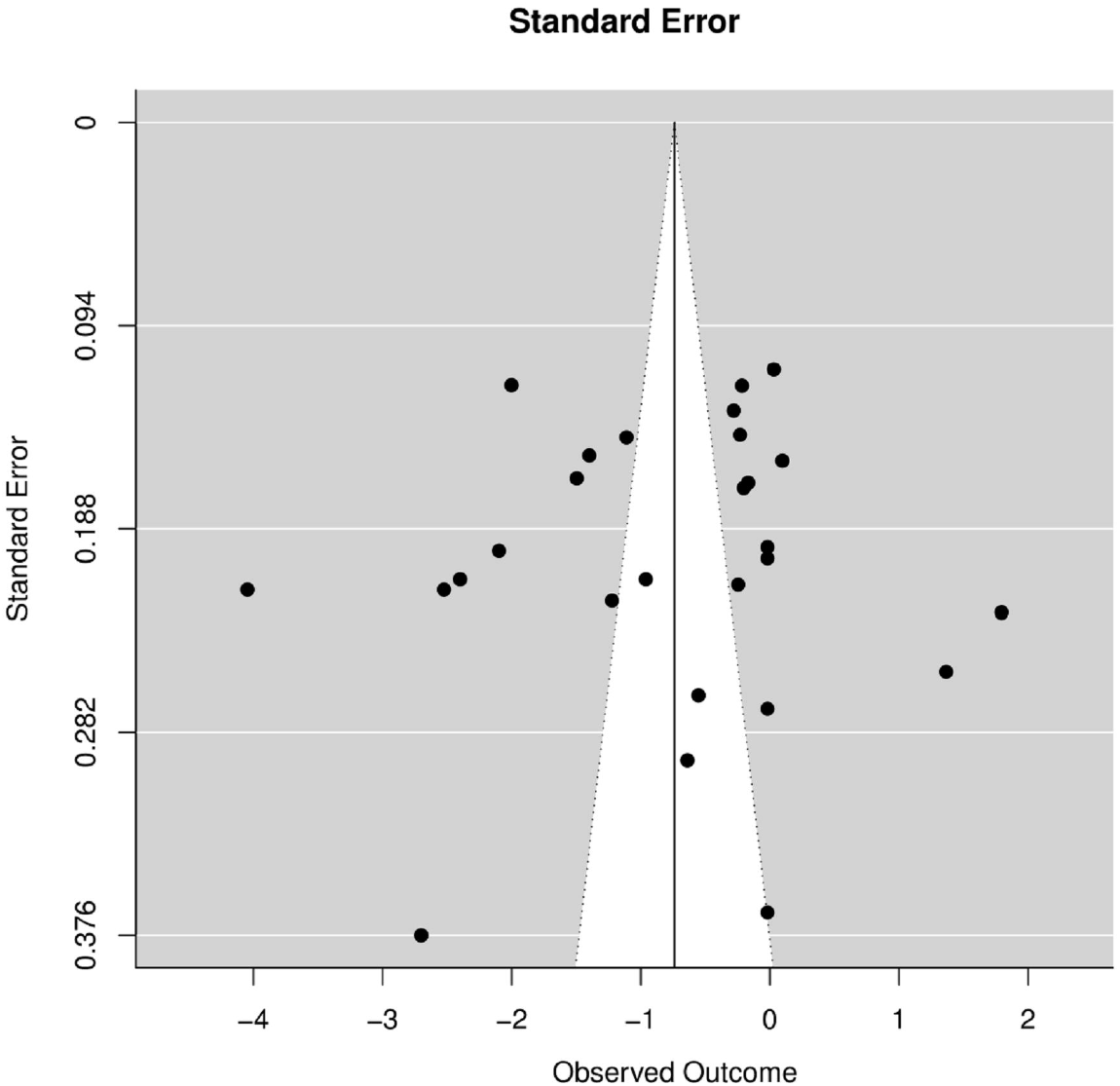
Funnel plot of surrogate effect size and surrogate standard error.

Among the 27 studies, 11 studies have primary goals to detect the AC-TL association. Among the 11 studies, two were cohort studies[16, 34], two were case-control studies[14, 15], and seven were cross-sectional studies[14, 16, 19, 25, 29, 34, 40]. The other 16 studies had various primary goals, from the association between TL and red blood cell size[24] to the association between TL and mortality in humans[38].

As for sex-specificity, eight studies are men only studies[14, 16, 20, 23, 27, 30], four studies are women only studies[19, 20, 29, 30], and 15 studies contain both men and women[15, 17, 18, 21, 22, 24-26, 28, 34, 38-40]. As for the categorization of AC, nine studies have the category of non-alcohol drinkers[14, 15, 17, 18, 22, 23, 25, 40], while two studies did not provide this information[20]. As for study population, 16 studies were based on populations in Europe[14, 16, 20-23, 26, 30, 34, 38], six studies were based on populations in America[19, 24, 25, 27-29], four studies were based on Asian[15, 17, 18, 40], and one study was based on Australian[39]. The range of total number of samples among the 27 articles is from 50[15] to 5862[29]. The mean age is ranged from 35 year old[25] to 79 year old[21]. The p-value is ranged from 9.00E-5 to 0.962.

Ten out of the 27 studies reported significant associations between AC and TL (i.e., p-value < 0.05). The Fisher’s combined p-value of the 27 studies is 5.75E-8 and the Liptak’s combined p-value is 8.76E-3.

Two out of the three cohort studies and all four case-control studies reported significant AC-TL associations, while only four out of the 20 (20%) cross-sectional studies reported significant AC-TL associations. For the association between study type and AC-TL association, the cross-table is in **S2 Table** online and the parallel pie chart is in **Fig 3.** Fisher’s exact tests showed that study type is significantly associated with the significance of AC-TL association (p-value=1.86E-3).

**Fig 3.**
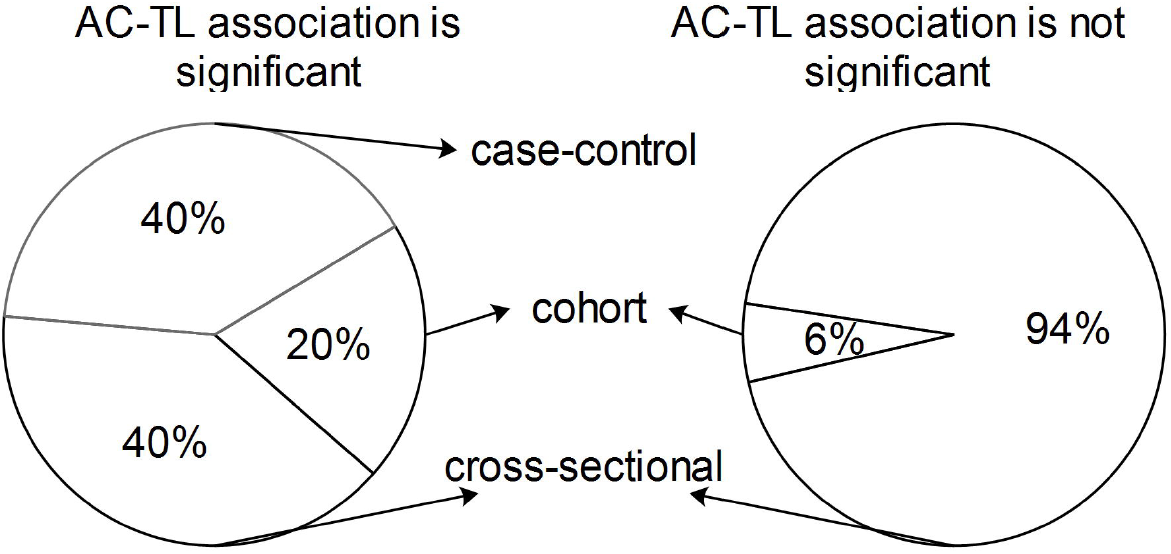
Parallel pie chart: The association between study type and AC-TL association.

All six studies from America reported non-significance of the AC-TL association, while all four studies from Asia and the only one study from Australia reported significant AC-TL association. Five out of 16 studies from Europe reported significant AC-TL association. For the association between continent and AC-TL association, the cross-table is in **S3 Table** online and the parallel pie chart is in **Fig 4.** Fisher’s exact test showed that study population (p-value = 2.67E-3) is significantly associated with the significance of the AC-TL association.

**Fig 4.**
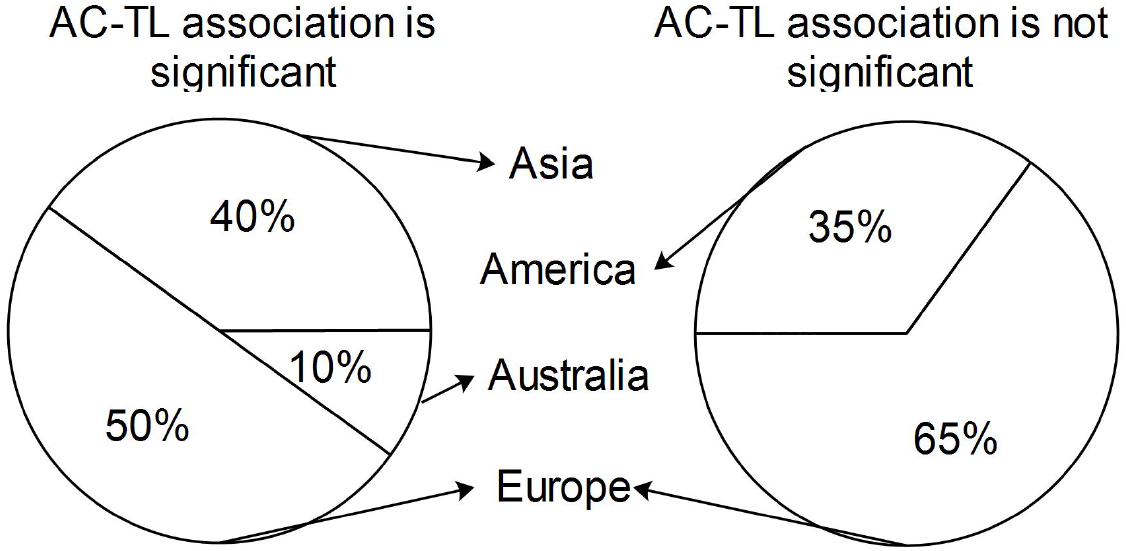
Parallel pie chart: The association between continent and AC-TL association.

The significance of AC-TL association is not related to whether a study’s primary goal is to test for AC-TL association (p-value=0.22), whether a study is based only on men, only on women, or on both men and women (p-value=0.39), or whether a study has an AC category: never drinker (p-value=0.09). Cross-tables of the association between these factors and AC-TL association are in **S4-S6 Tables** online, respectively.

Wilcoxon rank sum tests showed that both the total number of sample size (p-value=0.51) and mean age (p-value=0.11) are not associated with the significance of AC-TL association. Parallel boxplots of the association between these factors and AC-TL association are in **S1** and **S2 Figs** online.

## Discussion

In this article, we conducted a meta-analysis to summarize the pieces of evidence in the literature of observational studies about the association between alcohol consumption (AC) and telomere length (TL), and to identify factors that might affect the significance of the AC-TL association. There are 21 eligible articles included in this meta-analysis, containing 44 studies of the AC-TL association. The pooled evidence from the 27 independent studies showed that AC is significantly associated with TL (Fisher’s combined p-value=3.52E-8; Liptak’s combined p-value=8.24E-3). Study type (p-value=1.86E-3) and study population (p-value=2.67E-3) are significantly associated with the significance of the AC-TL association.

Using all 44 studies and ignoring the dependency among the 44 studies, The AC-TL association still remained significant (Fisher’s combined p-value=1.11E-16 and Liptak’s combined p-value=3.57E-5).

If we considered only the four case-control studies and the 20 cross-sectional studies, whether or not having an AC category “never drinker” is significantly associated with the significance of the AC-TL association (p-value=0.026). All four case-control studies reported significant AC-TL associations, while only four out of 20 cross-sectional studies reported significant AC-TL association. By observing that the controls in all four case-control studies are never drinkers and that only five out of the 20 cross-sectional studies had the never-drinker category, we guess that the effect of study type on the significance of the AC-TL association is probably due to the comparison of ever drinkers to never drinkers. Parallel pie chart and the cross-table of the association between whether a study has an AC category as never-drinker and the AC-TL association are in **S3 Fig** and **S7 Table** online.

Since there is only one study from Australia, we regroup study population to four Asia studies and 23 non-Asia studies; the study population is still significantly associated with the significance of the AC-TL association (p-value=0.012). Parallel pie chart and the cross-table of the association between whether the population of a study was Asian and the AC-TL association are in **S4 Fig** and **S8 Table** online. It would be interesting to investigate why all four studies from Asia reported significant AC-TL association.

It is interesting to observe that all four women-only studies reported non-significant associations between AC-TL, although sex-specificity is not significantly associated with the significance of the AC-TL association (p-value=0.3948). More women-only studies are needed to confirm if this observation is by chance or not.

Theoretically, longitudinal study (cohort study) would be better than cross-sectional study or case-control study in that cohort study could infer if heavy alcohol consumption causes shorter telomere length. However, there are several challenges for cohort studies[38]. First, it would be hard to make sure the procedure to measure TL at the end of long follow-up is the same as that at the baseline. Second, the blood storage method would also be different between baseline and the end of follow-up. Third, it would be difficult to make sure TL technically behaves in the same manner at baseline as that at the end of follow-up.

Genetics may play an important role in the AC-TL association. Pavanello et al. (2011) showed that carriers of the common ADH1B*1/*1 (rs1229984) genotype were more likely to be alcohol abusers, while exhibiting shorter TL[14]. Shin et al. (2016) showed that heavy alcohol consumption was inversely associated with leukocyte TL only among carriers of the mutant alleles (CT and TT) of rs2074356 (ALDH2)[40]. These studies indicate genetics could help identify subtypes of subjects that have significant AC-TL associations.

Several studies only provided lower or upper boundaries of p-values. For instance, when evaluating the effect of interaction between AC and ADH1B genotypes on TL among 255 controls, Pavanello et al. (2011) showed that the p-value < 0.001[14]. Pavanello et al. (2011) showed p-values < 0.0001 for another four studies[14]. Harris et al. (2006) showed p-value > 0.05[21]. When we calculated Fisher’s combined p-value and Liptak’s weighted p-value, we set p-value = 0.0009 for the study with p-value < 0.001, set p-value =0.00009 for the four studies with p-value < 0.0001, and set p-value = 0.962 (the maximum p-value among the 44 studies) for the study with p-value > 0.05.

Several studies provided mean age only for each group of subjects (e.g., for each TL tertile in Houben et al. (2010)[23]). We took the average of the means of these groups when we investigated if age would affect the significance of the AC-TL association.

Six of the 21 articles included more than one study that investigated the association between AC and TL. Among the 27 studies, two studies (one is men only study and the other is women-only study) were from Bekaert et al. (2007)[30]. The same is for Fyhrquist et al. (2011)[20]. Pavanello et al. (2011) included two studies: one is case-control study, and the other is cross-sectional study[14]. Stranberg et al. (2012) and Weischer (2014) both included two studies: one is cross-sectional study, and the other is cohort study[16, 34]. We assumed that the significance of the AC-TL association in different types of studies within the same article is independent.

The quantifications of TL are quite different among the 27 analyses. Sixteen analyses quantified TL as proportion or ratio. Among the 16 analyses, one study used normalized telomere-to-centromere ratio (NTCR)[15] and fifteen studies used the relative telomere to single copy gene (T/S) ratio[14, 17-19, 22, 25, 27, 28, 34, 38-40]. One study assessed TL with Z-score of T/S ratio[29]. One study assessed TL by terminal restriction fragment (TRF)[26]. Nine analyses used kilobase (kb) as the unit of TL. Three analyses divided TL into three[23, 24] or four categories[34]. Three analyses took the logarithm of TL[20, 40]. The difference in the TL quantification makes it difficult to aggregate data and interpret results. In future, it would be beneficial to have guidance on how to standardize the quantification of TL.

The quantifications of AC are also quite different among the 27 analyses. Twenty-three analyses measured AC on a continuous scale as alcoholic beverages per week[17, 21, 25, 28, 30, 38, 39], beverages per day[14, 29, 34], grams (g) per week[16], and g per day[18, 19, 23, 24, 26, 27, 40]. Among them, a standard alcoholic beverage widely varied from 12 oz. beer (1 bottle), 5 oz. wine (1 glass), 1.5 oz. liquor, to 12 g alcohol. One analysis measured AC as total drink-years (yearly frequency * total years of alcohol use)[22]. Other studies used categorized AC. One defined an ever-drinker as a person drinking at least one serving of beer (12 oz.), wine (4 oz.), or liquor (1.5 oz.) per month for ≥6 months, while a study used a four-category AC categorization: nondrinker, <10, 10-29, and >29 years of drinking alcohol[22]. Two analyses took the natural logarithm of AC[30]. Three analyses did not report the quantifications of AC[15, 20]. In future, it would be beneficial to have guidance on how to standardize the quantification of AC.

Among the 27 main studies investigated in the present study, 15 studies reported test statistic values, four of which reported positive associations between AC and TL[17, 18, 27, 29]. That is, the more alcohol consumption is, the longer telomere length is. Two of the four positive associations are statistically significant[17, 18]. Liu et al. (2009)[17] gave a potential explanation the significant positive association: telomerase activation by alcohol drinking in target tissues. They also mentioned that they could not rule out the possibility of chance findings because of the limited size in each subgroup[17]. We hypothesize that the positive AC-TL association is due to moderate drinking, which is beneficial to human health. However, Liu et al. (2009) did not mention how they defined “ever drinkers”[17]. Further investigation is warranted to study the differences of the AC-TL association among never drinkers, moderate drinkers, and heavy drinkers.

In summary, the present meta-analysis aggregated the p-values of 27 related studies (total sample size 35,891) and showed a significant association between alcohol consumption and telomere length based on Fisher’s combined p-value and Liptak’s weighted p-value, although the different quantification methods of AC and TL hindered the use of meta-analysis to aggregate the evidence for the AC-TL association. Factors, such as study type (case-control, cross-sectional, or cohort study), study population (American, Asian, or European), and subject gender, might affect the AC-TL association. In future, it would help us uncover the true relationship between alcohol consumption and telomere length if we standardize the quantifications of AC and TL and if we take account for both beneficial and deteriorating effects of AC to human health.

## Supporting information

**S1 Fig. Parallel boxplots of the association between total number of sample size and AC-TL association.**

(PDF)

**S2 Fig. Parallel boxplots of the association between mean age and AC-TL association.**

(PDF)

**S3 Fig. Parallel pie chart: The association between whether a study has an AC category as never-drinker and the AC-TL association (no cohort).**

(PDF)

**S4 Fig. Parallel pie chart: The association between whether the population of a study was Asian and the AC-TL association**

(PDF)

**S1 Checklist. PRISMA checklist.**

(DOC)

**S1 Table. The information of the 44 analyses in the 21 articles.**

(xlsx)

**S2 Table. Cross table: The relationship between study type and the significance of the AC-TL association. Note: ratio: the ratio of number of studies with significant AC-TL association to the number of total studies**

(xlsx)

**S3 Table. Cross table: The relationship between continent and the significance of the AC-TL association.**

(xlsx)

**S4 Table. Cross table: The relationship between a study’s primary goal and the significance of the AC-TL association.**

(xlsx)

**S5 Table. Cross table: The relationship between gender of a study’s participants and the significance of the AC-TL association.**

(xlsx)

**S6 Table. Cross table: The relationship between whether a study has an AC category as never-drinker and the significance of the AC-TL association.**

(xlsx)

**S7 Table. Cross table: The relationship between whether a study has an AC category as never-drinker and the significance of the AC-TL association (cohort study excluded).**

(xlsx)

**S8 Table. Cross table: The relationship between whether the population of a study was Asian and the significance of the AC-TL association.**

(xlsx)

## Author Contributions

**Conceptualization:** Jianqiang Li, Ji-Jiang Yang, Weiliang Qiu.

**Data curation:** Yu Guan, Faheem Akhtar, Xi Xu.

**Investigation:** Yu Guan, Jianqiang Li, Weiliang Qiu.

**Methodology:** Jianqiang Li, Ji-Jiang Yang, Weiliang Qiu.

**Project administration:** Jianqiang Li, Ji-Jiang Yang.

**Software:** Yu Guan, Faheem Akhtar, Xi Xu.

**Supervision:** Shi Chen, Qing Wang, Hui Pan.

**Validation:** Yu Guan, Weiliang Qiu.

**Writing - original draft:** Faheem Akhtar, Xi Xu, Shi Chen, Qing Wang, Hui Pan.

**Writing - review & editing:** Shi Chen, Qing Wang, Hui Pan.

